# A novel USP30 inhibitor recapitulates genetic loss of USP30 and sets the trigger for PINK1-PARKIN amplification of mitochondrial ubiquitylation

**DOI:** 10.1101/2020.04.16.044206

**Authors:** Emma Rusilowicz-Jones, Jane Jardine, Andreas Kallinos, Adan Pinto-Fernandez, Franziska Guenther, Mariacarmela Giurrandino, Francesco G. Barone, Katy McCarron, Christopher J. Burke, Alejandro Murad, Aitor Martinez, Elena Marcassa, Malte Gersch, Alex Buckmelter, Katherine J. Kayser-Bricker, Frederic Lamoliatte, Akshada Gajbhiye, Simon Davis, Hannah C. Scott, Emma Murphy, Katherine England, Heather Mortiboys, David Komander, Matthias Trost, Benedikt M. Kessler, Stephanos Ioannidis, Michael Ahlijanian, Sylvie Urbé, Michael J. Clague

## Abstract

The mitochondrial deubiquitylase USP30 negatively regulates the selective autophagy of damaged mitochondria. It has been proposed as an actionable target to alleviate the loss of function of the mitophagy pathway governed by the Parkinson’s Disease associated genes PINK1 and PRKN. We present the characterisation of a N-cyano pyrrolidine derived compound, FT3967385, with high selectivity for USP30. The compound is well tolerated with no loss of total mitochondrial mass. We demonstrate that ubiquitylation of TOM20, a component of the outer mitochondrial membrane import machinery that directly interacts with USP30, represents a robust biomarker for both USP30 loss and inhibition. We have conducted proteomics analyses on a SHSY5Y neuroblastoma cell line model to directly compare the effects of genetic loss of USP30 with selective inhibition in an unbiased fashion. We have thereby identified a subset of ubiquitylation events consequent to mitochondrial depolarisation that are USP30 sensitive. Within responsive elements of the ubiquitylome, several components of the outer mitochondrial membrane transport (TOM) complex are most prominent. Thus, our data support a model whereby USP30 can regulate the availability of ubiquitin at the specific site of mitochondrial PINK1 accumulation following membrane depolarisation. In this model, USP30 deubiquitylation of TOM complex components dampens the trigger for the Parkin-dependent amplification of mitochondrial ubiquitylation leading to mitophagy. Accordingly, PINK1 generation of phospho-Ser65 Ubiquitin proceeds more rapidly and to a greater extent in cells either lacking USP30 or subject to USP30 inhibition.

## Introduction

Damaged mitochondria are removed from the cell by a process of selective autophagy termed mitophagy. Defects in mitochondrial turnover have been linked to a number of neurodegenerative conditions including Parkinson’s Disease (PD), Alzheimer’s Disease (AD) and Motor Neuron Disease (MND) (1, 2). This process is best understood in the context of PD, for which loss of function mutations in the mitophagy promoting genes *PINK1* and *PRKN* (coding for the Parkin protein) are evident (3, 4). Mitochondrial depolarisation leads to the accumulation of the PINK1 kinase at the mitochondrial surface, which then phosphorylates available ubiquitin moieties at Ser65 (5-9). PhosphoSer65-Ubiquitin (pUb) recruits the ubiquitin E3 ligase Parkin to mitochondria, where it is fully activated by direct PINK1-dependent phosphorylation at Ser65 of its UBL domain (10-13). This triggers a feed-forward mechanism that coats mitochondria with ubiquitin, leading to selective engulfment by autophagosomal membranes (14, 15).

The deubiquitylase (DUB) family of enzymes plays a role in most ubiquitin dependent processes, by promoting ubiquitin flux or suppressing ubiquitylation of specific substrates (16, 17). USP30 is one of only two DUBs that possess a transmembrane domain. Its localisation is restricted to the outer mitochondrial membrane and to peroxisomes (18-21). USP30 can limit the Parkin-dependent ubiquitylation of selected substrates and depolarisation-induced mitophagy in cell systems that have been engineered to over-express Parkin (22-25). We have recently shown that it can also suppress a PINK1-dependent component of basal mitophagy, even in cells that do not express Parkin (20). Thus USP30 may represent an actionable drug target relevant to PD progression and other pathologies to which defective mitophagy can contribute (26-28). One attractive feature of USP30 as a drug target in this context, is that its loss is well tolerated across a wide range of cell lines (29).

The Ubiquitin Specific Protease (USPs) DUB family are cysteine proteases and comprise around 50 members in humans (17). Early academic efforts to obtain specific small molecule inhibitors were only partially successful (30). More recently industry-led efforts have generated some highly specific inhibitors, exemplified by compounds targeting USP7, an enzyme linked to the p53/MDM2 signaling axis (31-35). Some N-cyano pyrrolidines, which resemble known cathepsin C covalent inhibitors, have been reported in the patent literature to be dual inhibitors of UCHL1 and USP30 (36). High-throughput screening has also identified a racemic phenylalanine derivative as a USP30 inhibitor (37). However the specificity and biological activity of this compound has so far been only characterised superficially.

Here we introduce FT3967385 (hereafter FT385), a modified N-cyano pyrrolidine tool compound USP30 inhibitor. We carefully correlate its effects upon the proteome and ubiquitylome of neuroblastoma SH5YSY cells, expressing endogenous Parkin. We also show that this compound can recapitulate effects of USP30 deletion on mitophagy and regulate the ubiquitin status of Translocase of the Outer Mitochondrial Membrane (TOM) complex components. The TOM complex functions as a common entry portal for mitochondrial precursor proteins (38). We propose that associated ubiquitin may provide nucleating sites at which PINK1 phosphorylation sets in train a feed-forward loop of further Parkin-mediated ubiquitylation (24). Accordingly, pUb generation following mitochondrial depolarisation is enhanced by both USP30 deletion and by inhibitor treatment.

## Results

We developed a tool compound inhibitor (FT385) for investigation of USP30 biology (Figure 1A). It shows a calculated IC_50_ of ~1nM *in vitro* using purified USP30, together with ubiquitin-rhodamine as a fluorogenic substrate (Figure 1B,E). Bio-layer interferometry experiments show binding behaviour that is consistent with covalent modification of USP30 (Figure 1C). To test for selectivity of the inhibitor within the USP family of enzymes, we used the Ubiquigent DUB profiler screen, which tests inhibitory activity against a broad panel of USP enzymes. At the indicated concentrations (up to 200nM) the inhibitor was highly selective for USP30 (Figure 1F). Only one other family member, the plasma membrane associated USP6, showed a significant degree of inhibition (19). This particular deubiquitylase shows a highly restricted expression profile (39).

**Figure 1.**
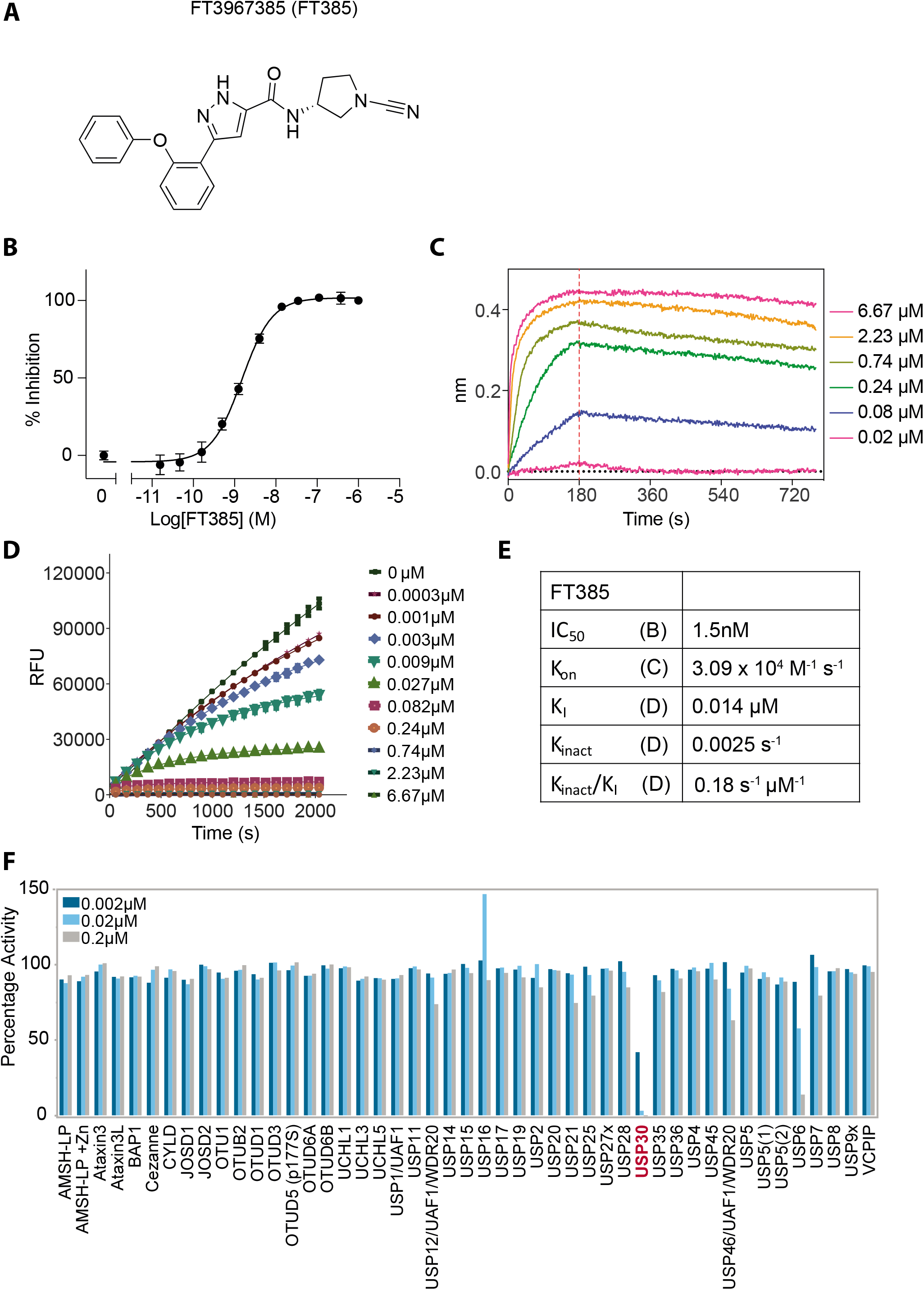
FT3967385 is a selective covalent USP30 inhibitor. (A) Chemical structure of FT3967385 (FT385). (B) Concentration dependent inhibition of recombinant USP30 activity using ubiquitìn-rhodamine as a substrate. (C) Bio-layer interferometry (BLI) traces showing no significant off-rate at indicated concentratìons.Red line indicates removal of the inhibitor after 180 seconds. (D) Progress curves characteristic of a covalent inhibitor (0-6.67μM), these are fitted to obtain K_I_ and K_inact_. (E) Data table of inhibitory properties. (F) DUB specificity screen (DUB profiler, Ubiquigent) with 2, 20 and 200nM FT385.

We used the competition between FT385 and Ub-propargylamide (Ub-PA), which covalently binds to the USP30 active site, to assess target engagement (40). Binding of the probe to a DUB leads to an up-shift in apparent molecular weight on SDS-PAGE gels (Figure 2). If a drug is present that occupies or otherwise occludes this site, probe modification is inhibited and the protein mass is down-shifted accordingly. Our results demonstrate target engagement and allow us to determine a suitable concentration range for further experiments (Figure 2). In SHSY5Y neuroblastoma cells, effective competition of drug towards added probe is seen at concentrations >100nM when added to cell lysates (Figure 2A) or pre-incubated with cells prior to lysis (Figure 2B).

**Figure 2.**
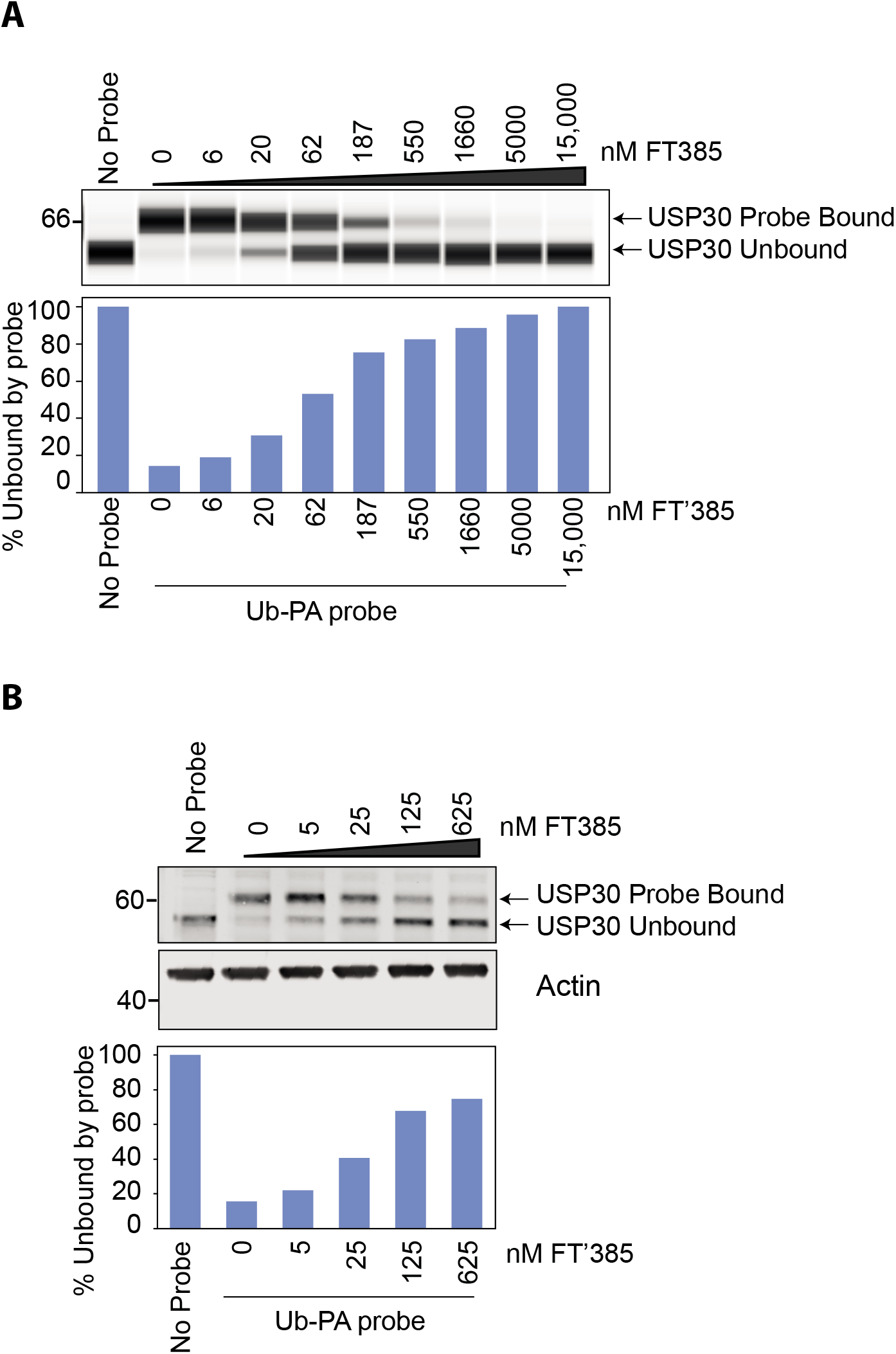
Activity based ubiquitin probe assay shows that FT385 engages USP30 in cells at low nanomolar concentrations. (A) SHSY5Y cell homogenates or (B) intact SHSY5Y cells were incubated with FT385 for 30 minutes or 4 hours respectively at the indicated concentrations, then incubated with Ub-PA probe for 15 minutes at 37°C and immunoblotted as shown. Samples in (A) were analysed using an automated western blot (WES^™^) system.

To be able to compare compound activity to USP30 loss, we used CRISPR/Cas9 to generate YFP-Parkin-RPE1 (retinal pigment epithelium) and SH5Y5Y (neuroblastoma) USP30 knock-out (KO) cells (Supplementary Figure 1). We have previously shown that USP30 physically interacts with TOM20, a component of the outer mitochondrial membrane transport complex which recognises mitochondrial targeting sequences (24, 38). USP30 represses both depolarisation induced mitophagy and the specific ubiquitylation of TOM20 in cells over-expressing Parkin (22-24, 41). Application of FT385 to RPE1 cells over-expressing YFP-Parkin results in enhanced ubiquitylation and degradation of TOM20 without affecting PINK1 protein levels (Figure 3A). Enhancement of TOM20 ubiquitylation by FT385 under depolarising conditions is more clearly shown in Figure 3B. In this experiment, a shorter depolarisation time (1 hour) has been used, at which there is minimal TOM20 loss to mitophagy or other pathways. USP30 KO and inhibitor treated cells show similar elevation of ubiquitylated TOM20, whilst no further enhancement is achieved by inhibitor treatment of KO cells (Figure 3B). Thus, the TOM20 ubiquitylation response depends on USP30 catalytic activity and represents an on-target effect of the drug.

**Figure 3.**
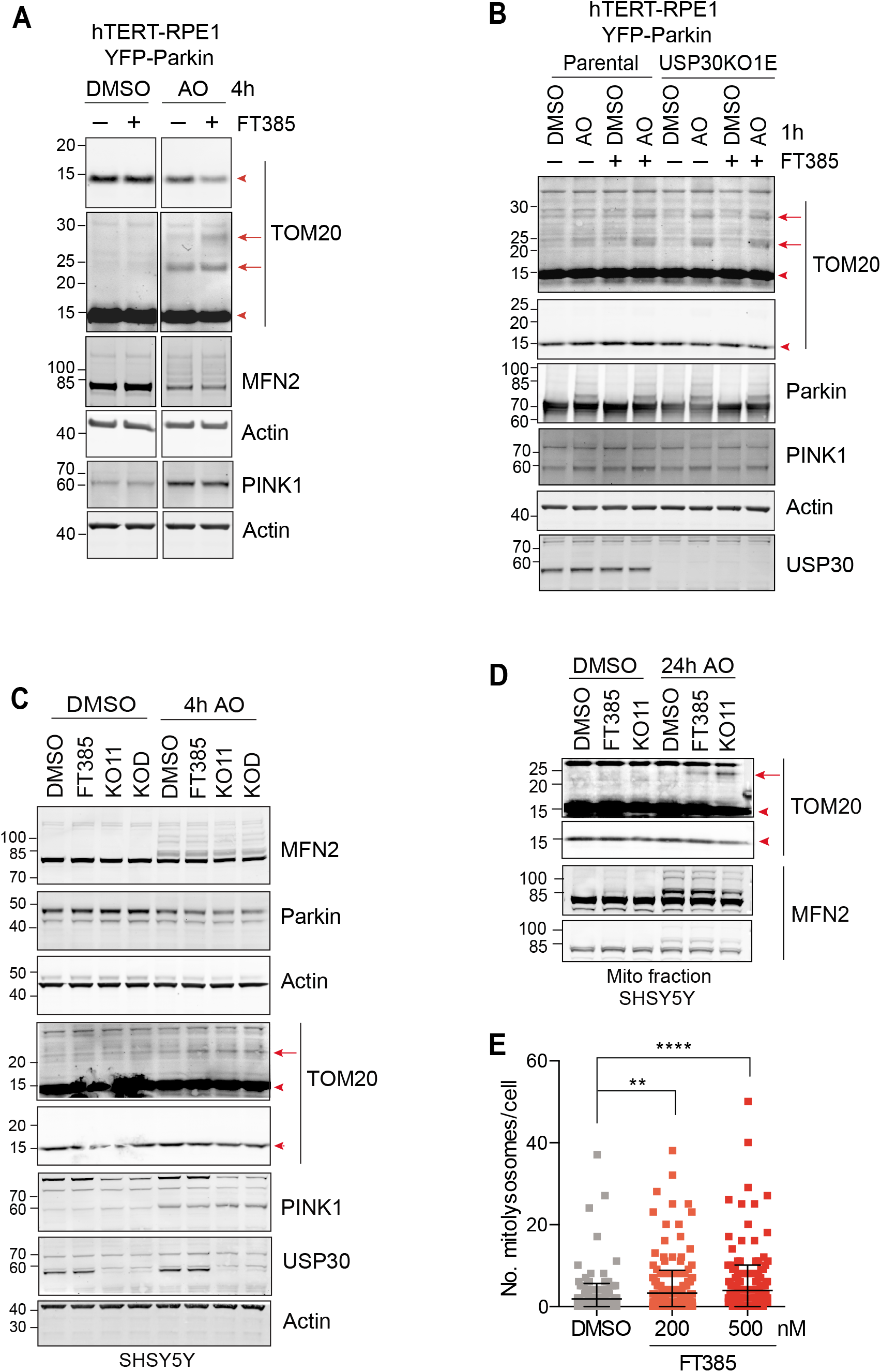
Pharmacological inhibition of USP30 phenocopies USP30 KO in enhancing basal mitophagy and promoting ubiquitylation of TOM20 upon depolarisation. (A) Inhibition of USP30 enhances the ubiquitylation and degradation of TOM20 in YFP-Parkin over-expressing hTERT-RPE1 cells in response to mitophagy induction. Cells were treated for 4 hours with DMSO or Antìmycin A and Oligomycin A (AO; 1μM each) in the absence or presence of 200 nM FT385, lysed and analysed by western blodng. Red arrows indicate ubiquitylated TOM20 and arrowheads indicate unmodified TOM20. (B) USP30 inhibitor (FT385) treatment of parental (PAR) YFP-Parkin overexpressing hTERT-RPE1 cells phenocopies USP30 deletion (KO1E) by promoting TOM20 ubiquitylation. In contrast, TOM20 ubiquitylation is unaffected by FT385 in the corresponding USP30 KO (KO1E) cells. Cells were treated for 1 hour with or without AO (1μM) in the absence or presence of 200 nM FT385, lysed and samples analysed by immunoblodng. (C) TOM20 ubiquitylation is enhanced by USP30 inhibition and deletion in SHSY5Y cells expressing endogenous Parkin levels. SHSY5Y with or without FT385 (200nM) and USP30 CRISPR/Cas9 KO cells (KO11 and KOD, two distinct sgRNAs) were treated with AO (1μM each) for 4 hours as indicated. Cells were then lysed and samples analysed by immunoblodng as shown. (D) SHSY5Y-PAR (mitoQC) and USP30 KO cells (KO11) were treated for 24 hours with AO (1μM each) in the presence or absence of FT385 (100nM). Cells were subjected to sub-cellular fractionation and the mitochondria-enriched fraction (MF) was analysed by immunoblodng as indicated. (E) Quantification of the number of mitolysosomes in SH-SY5Y-mitoQC cells, treated with DMSO or FT385 (200 or 500nM) for 96 hours prior to imaging. Average ± SD; *n*=3 independent experiments; 80 cells per experiment; one-way ANOVA with Dunnett’s multiple comparisons test, *\*\*P* < 0.01, \*\*\*\**P* < 0.0001.

We confirmed that both USP30 deletion and inhibition can also lead to the accumulation of ubiquitylated TOM20 in SHSY5Y cells, both in whole cell lysates and in crude mitochondrial fractions (Figure 3C,D). Notably, this is the first time that this modification has been detected by Western blotting without Parkin over-expression. TOM20 is atypical in the respect that we do not observe USP30-dependent changes to the ubiquitylation pattern of another mitochondrial Parkin substrate Mitofusin 2 (MFN2) (Figure 3A,C,D). To determine effects of USP30 inhibition on basal mitophagy, we used SHSY5Y expressing a tandem mCherry–GFP tag attached to the outer mitochondrial membrane localization signal of the protein FIS1 (42). A clear increase in the number of mitolysosomes per cell, indicative of increased mitophagic flux, is apparent following USP30 inhibition over a 96 hour time period (Figure 3E).

Trypsin digestion of ubiquitylated proteins generates peptides with a residual diGly motif, which provides a characteristic mass shift and can be used for enrichment by immunoprecipitation (43). Several studies have used this approach to define Parkin substrates through proteomic analysis, following mitochondrial depolarisation in cell lines over-expressing Parkin (8, 44, 45). In order to search for potential substrates and/ or biomarkers beyond TOM20, we decided to take an unbiased view of USP30 control of the cellular proteome and ubiquitylome, in SHSY5Y cells, which endogenously express Parkin. Our experimental design, using triplexed combinations of SILAC labels, allowed quantitative comparison of both USP30 inhibitor treated cells (200nM) and USP30 KO cells relative to parental untreated cells in basal conditions (proteome) or following mitochondrial depolarisation (proteome + ubiquitylome) (Figure 4A). We quantitated 6,562 proteins and 9,536 diGly peptides (which indicate specific sites of ubiquitylation), derived from 2,915 proteins (Supplementary Table 1). We had hoped that the proteome might provide a biomarker that could be used in pre-clinical models for testing drug efficacy. Despite obtaining deep proteome coverage, we identified few proteins that responded to both genetic deletion and inhibition of USP30 (24 hours) in a consistent manner across experiments. No impact of USP30 on total mitochondrial or peroxisomal mass following 24 hours depolarisation is apparent (Figure 4B). This is in keeping with our observations and previous findings, that in cell lines expressing endogenous levels of Parkin, the extent of depolarisation-induced mitophagy is low (46). In this experiment we find that USP30 influences the ubiquitylation status of a small minority of proteins following depolarisation (Figure 4C). Most prominent among them are members of the voltage dependent anion channel (VDAC) family. VDAC1, VDAC2 and VDAC3 show enhanced ubiquitylation at specific sites in the absence of USP30 activity without any change at the proteome level. In general, the effect is stronger in the USP30 KO cells but the pattern is conserved with USP30 inhibitor treatment (Figure 4C-E, Supplementary Figure 2). One conclusion from these data is that the global impact of USP30 activity at both the proteome and ubiquitylome levels is subtle. This makes pharmacology experiments in both terminally differentiated cellular experiments (e.g., primary cultured rodent neurons or human iPSC-derived neurons) and *in vivo* experiments challenging. However, it is consistent with low impact on cell viability seen in CRISPR screens (47) and may in fact be a desirable feature of a drug target for a neurodegenerative disease.

**Figure 4.**
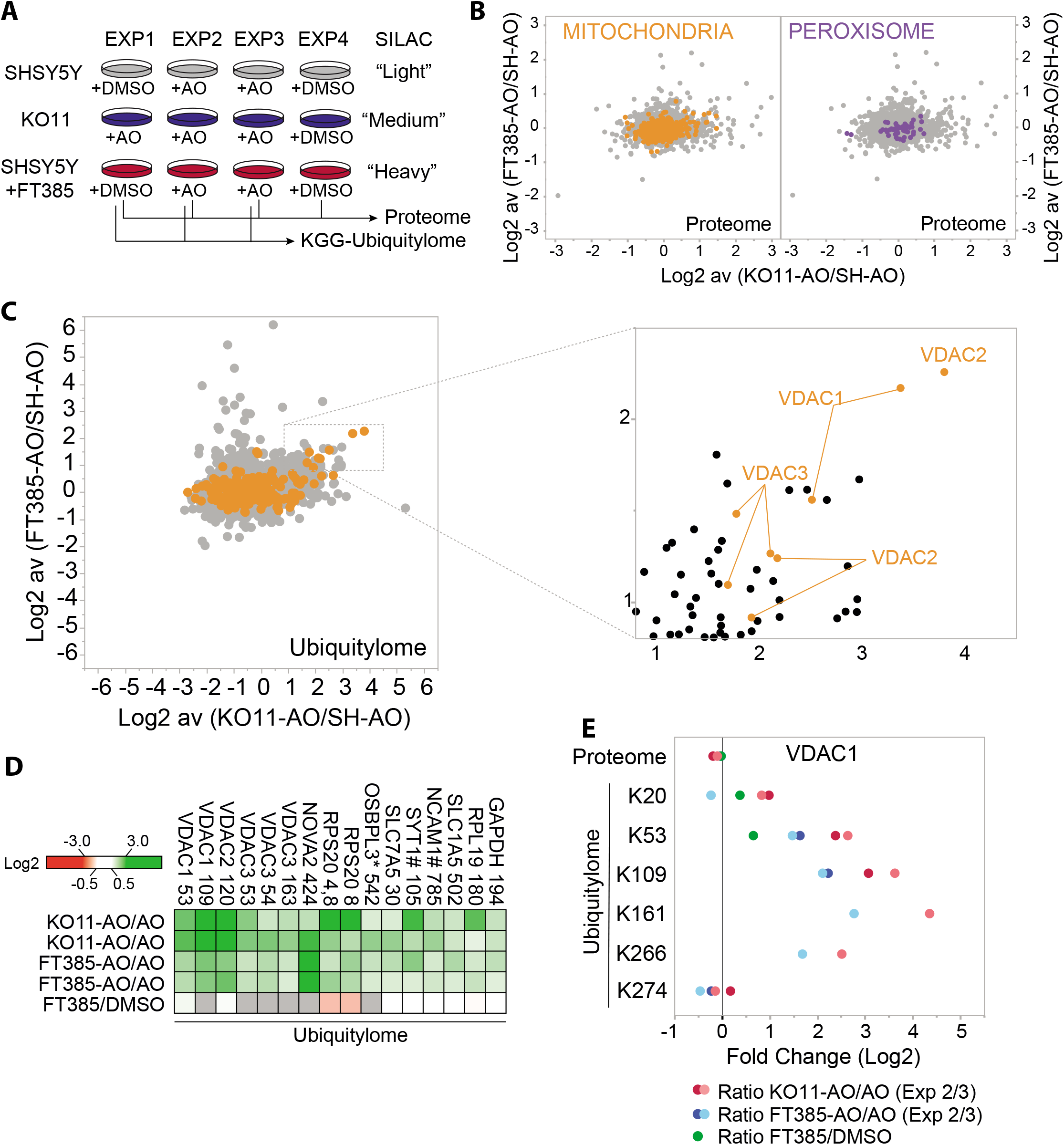
Comparison of Proteome and Ubiquitylome changes in USP30 KO versus USP30 inhibitor treated SHSY5Y cells. (A) Schematic flow-chart of SILAC based quantitative ubiquitylome and proteome analysis comparing USP30 KO and USP30 inhibition. SHSY5Y (USP30 wild-type) and SHSHSY USP30 KO (KO11) cells were metabolically labelled by SILAC as shown. Cells were then treated for 24 hours with DMSO or Antìmycin A and Oligomycin A (AO; 1μM each) and/or FT385 (200nM) as indicated. Cells were lysed and processed for mass spectrometry analysis. Graphs depicting the fold change (log2) in the proteome (B) or ubiquitylome (C) of AO-treated SHSY5Y cells ± FT385 treatment (y-axis) and ± USP30 (x-axis). Mitochondrial (Integrated Mitochondrial Protein Index (IMPI) database; http://www.mrc-mbu.cam.ac.uk/impi; “known mitochondrial” only) and peroxisomal proteins (peroxisomeDB; http://www.peroxisomedb.org) proteins are highlighted in orange and purple respectively. Inset in (C) shows enlarged section of ubiquitylome data for peptides enriched in USP30 KO and inhibitor treated cells. Within proteome graphs (B) each dot represents a single protein identified by at least 2 peptides and the ratio shows the average of 2 experiments. Within ubiquitylome graphs (C) each dot represents a single diGLY peptide (localisation ≥0.75) and the ratio shows the average of 2 experiments. (D) Heatmap showing diGly peptides that are increased consistently by log2 ≥ 0.8 in both USP30 KO and USP30 inhibitor (FT385) treated cells. * indicates ambiguity of peptide assignment between family members (OSBPL3,OSBPL7,OSBPL6). Grey indicates the protein was not seen in that condition. # indicates an increase at proteome level in KO11. VDAC3 K53 and K54 correspond to equivalent lysines in two distinct isoforms. (E) Fold change (Log2) in proteome and individual diGly peptides (localisation ≥0.75) by site in VDAC1 proteins. See Supplementary Figure 2 for corresponding datasets for VDAC2 and 3.

To obtain information on the early USP30-dependent changes to the mitochondrial ubiquitylation profile that follow depolarisation, we compared two USP30 KO SHSY5Y clones with wild type cells, using a shorter depolarisation period (4 hours, Figure 5A). No systematic changes in mitochondrial or peroxisomal protein abundance were observed (Figure 5B). For the ubiquitylome arm of this experiment we used crude mitochondrial fractions to increase coverage of specific mitochondrial components. This is evident in Figures 5C and 5D, which summarise the major changes in ubiquitylation we have identified at specific sites in both sets of experiments (Figures 4A and 5A and Supplementary Tables 1 and 2). Multiple responsive VDAC peptides were once again identified. Strong outliers are found in Ganglioside-induced differentiation associated protein 1 (GDAP1), an outer mitochondrial membrane protein, mutations of which are linked to Charcot-Marie-Tooth neuropathy and mitochondrial dysfunction (48) and the mitochondrial outer membrane protein Synaptojanin 2 binding protein (SYNJ2BP, Figure 5E) (49). Also prominent is Peptidyl-tRNA Hydrolase 2 (PTRH2), a mitochondrial protein linked to the release of non-ubiquitylated nascent chains from stalled ribosomal complexes (50). The improved coverage now reveals USP30-dependent ubiquitylation of multiple Tom complex components including the two translocase receptors, TOM20 and TOM70, the TOM40 channel and an accessory subunit TOM5 within this set of strong outliers.

**Figure 5.**
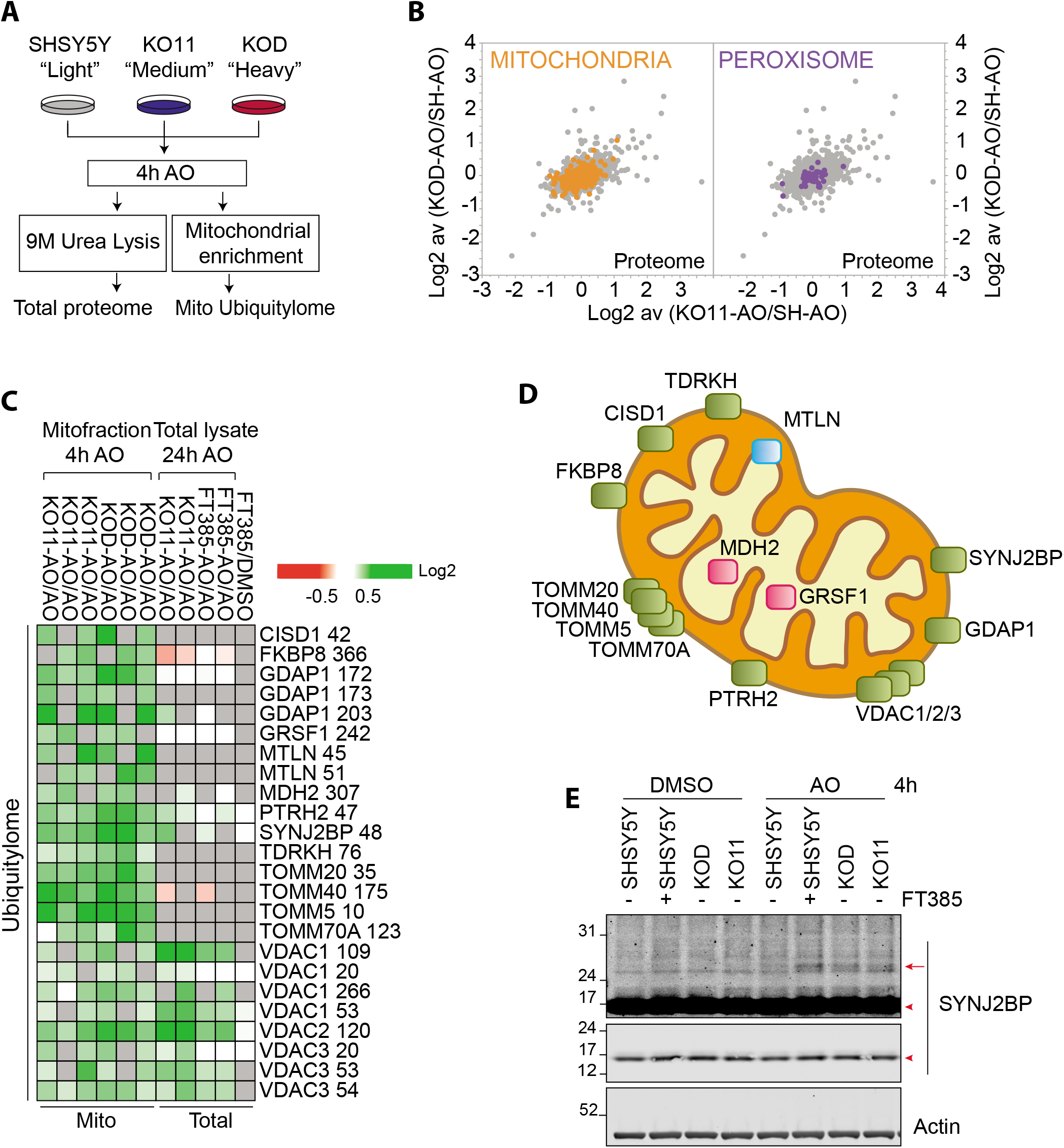
Proteomic analysis of the mitochondria-enriched ubiquitylome in USP30 KO SHSY5Y cells. (A) Schematic flow-chart of SILAC based quantitative ubiquitylome and proteome analysis comparing two USP30-KO clones (KOD-sgRNA#1; KO11-sgRNA#2) to wild-type SHSY5Y cells. Cells were metabolically labelled by SILAC as shown and treated for 4 hours with AO (1μM). Cells were then either lysed for total proteome analysis or further processed by subcellular fractionation. The mitochondrial fraction was used as the starting material for the ubiquitylome analysis. (B) Graphs depicting the fold change (log2) in the proteome of AO-treated USP30 KOD versus wildtype SHSY5Y (SH) (y-axis) and USP30 KO11 compared to SHSY5Y cells (x-axis). Mitochondrial (Integrated Mitochondrial Protein Index (IMPI) database; http://www.mrc-mbu.cam.ac.uk/impi; “known mitochondrial” only) and peroxisomal proteins (peroxisomeDB; http://www.peroxisomedb.org) proteins are highlighted in orange and purple respectively. Each dot represents a single protein identified by at least 2 peptides and the ratio shows the average of 3 experiments. (C) Heatmap showing diGLY containing peptides that are increased consistently in at least 4 out of 6 experimental conditions by log2 ≥ 0.8. The corresponding data from the total ubiquitylome experiment shown in Figure 4 are also indicated. Grey indicates the protein was not seen in that condition. VDAC3 K53 and K54 correspond to equivalent lysines in two distinct isoforms. (D) Depiction of the localisation of USP30 sensitive depolarisation-induced ubiquitylated proteins within mitochondria (enriched proteins shown in C). Defined as outer mitochondrial membrane (green), inner mitochondrial membrane (blue) or matrix (pink).(E) Western blot showing the appearance of mono-ubiquitylated species of SYNJ2BP in both USP30 KO clones (KO11, KOD) and in USP30 inhibitor (FT385) treated cells. Cells were treated for 4 hours with AO (1μM) in the presence or absence of 200nM FT385, then lysed in Urea lysis buffer and analysed by western blot. Red arrows indicate ubiquitylated SYNJ2BP and the arrowheads indicate unmodified SYNJ2BP.

In healthy mitochondria, PINK1 is imported through the TOM complex and subsequently cleaved and released for proteasomal degradation in the cytosol. In depolarised mitochondria it is no longer imported and degraded but remains associated with TOM complex components on the outer mitochondrial membrane (51-54). At this point it becomes trans-activated and initiates a signaling cascade by phosphorylating ubiquitin on Ser65 (generating pUb). This accumulation of pUb can be readily visualised by Western blotting using a specific antibody. We find that genetic loss of USP30 or USP30 inhibition both lead to a more rapid accumulation of pUb following mitochondrial depolarisation, without an evident increase in total PINK1 nor Parkin levels at mitochondria (Figures 6A-D and Supplementary Figure 3). The differential is more prominent at earlier time points following depolarisation, but elevated levels are sustained up to 24 hours (Figure 6C).

**Figure 6.**
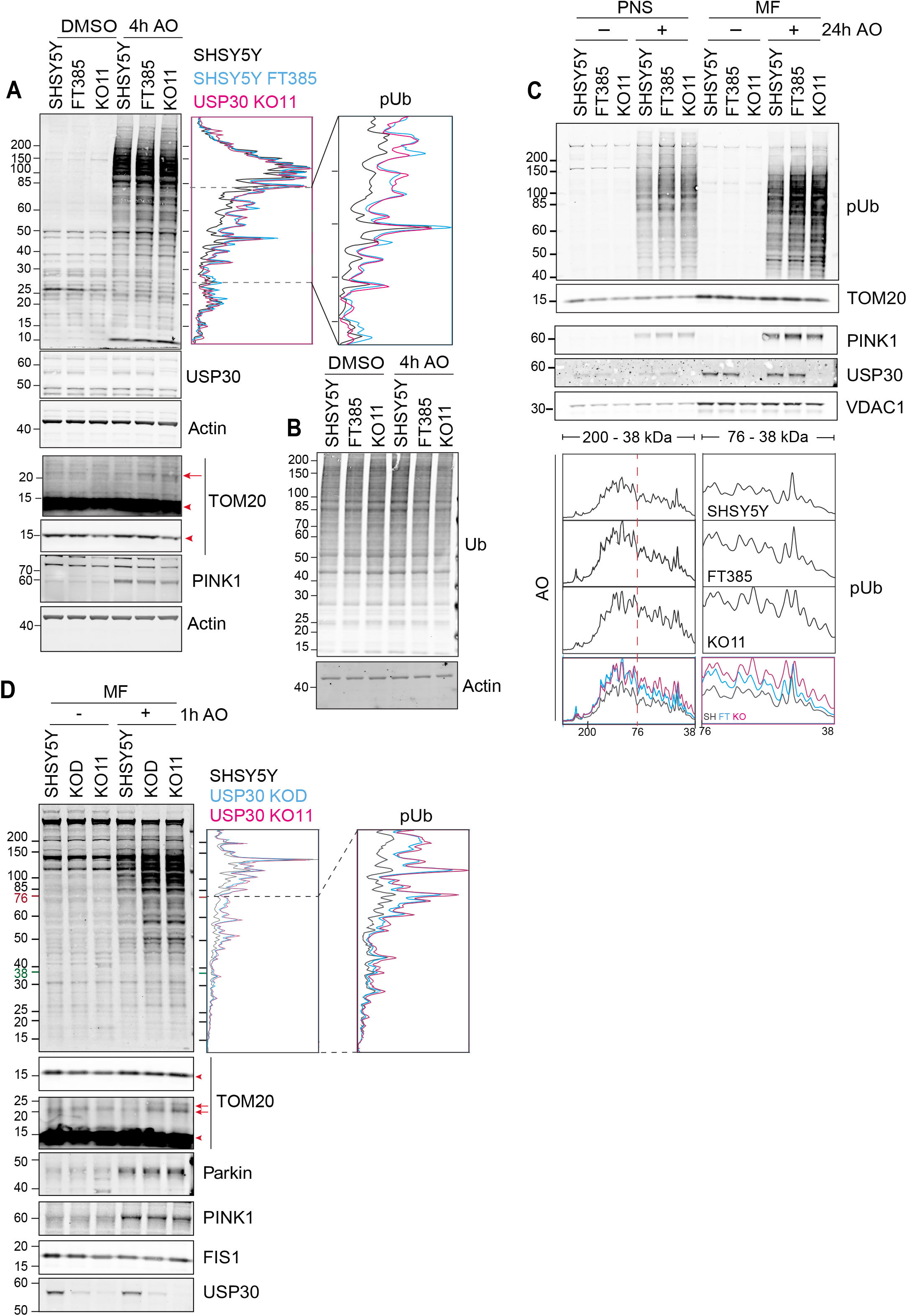
USP30 KO and USP30 Inhibition enhance phospho-Ser65 Ubiquitin levels on mitochondria of SHSY5Y cells. (A) Comparison of depolarisation induced phosphoSer65-Ubiquitìn (pUb) generation in SHSY5Y cells treated with FT385 and USP30 KO SHSY5Y (KO11). Shown is a western blot, and corresponding line-graph for the pUb signal, of lysates from cells treated for 4 hours with AO (1μM) with or without FT385 (200nM). Red arrows indicate ubiquitylated TOM20 and the arrowheads indicate unmodified TOM20. (B) Same samples as in (A) probed for total ubiquitin (VU1). (C) A post-nuclear supernatant (PNS) and mitochondrial fractions (MF) were obtained from SHSY5Y cells treated in presence or absence of FT385 (100nM, 24 hours), with DMSO or AO (1μM). Samples were analysed by western blodng and a line graph depicting the pUb signal are shown. (D) SHSY5Y cells and two USP30 KO clones (KOD and KO11) were treated for 1 hour with AO (1μM). Cells were homogenised and mitochondrial fractions (MF) prepared and analysed as indicated.

## Discussion

Here we provide a comprehensive analysis of the impact of USP30 on mitochondrial ubiquitylation dynamics following mitochondrial membrane depolarisation. Our principal analysis is conducted on cells expressing endogenous levels of Parkin and we directly compare the effects of genetic loss with a specific inhibitor. This allows us to clearly attribute molecular signatures to catalytic activity for the first time. We have extended USP30 linkage to the mitochondrial import (TOM) complex to now include sub-units beyond TOM20, which has been previously characterised (4, 24, 41). We also identify a further substrate, SYNJ2BP, whose enhanced ubiquitylation can be monitored by Western blotting. Based on our studies FT385 emerges as a promising tool compound for the study of USP30 biology. When used at appropriate concentrations, a high degree of specificity amongst DUB family members can be achieved. On the other hand, there are some inevitable liabilities; following inhibitor treatment, we identify several proteins with enhanced ubiquitylation that is not evident with genetic loss of USP30.

Previous studies have suggested that the overall pattern of depolarisation-induced ubiquitylation of mitochondria is largely unchanged following USP30 knockdown, with TOM20 being an exception (24, 41). We see enhanced pUb accumulation in the absence of USP30 activity, despite the published observations that pUb modified chains provide a poor substrate for USP30 (9, 41). How then might USP30 suppress mitophagy, as previously reported in several studies (20, 22-24)? We have previously shown that USP30 depletion enhances PINK1-dependent basal mitophagy even in the absence of Parkin (20). We and others have proposed that USP30 may regulate the availability of ubiquitin on specific trigger proteins that are most readily available for phosphorylation by PINK1. In other words, USP30 may determine the probability that a local accumulation of PINK1 can trigger feed-forward mechanisms that lead to mitophagy (20, 41, 55). The prominence of TOM complex components within the limited set of USP30-responsive diGly-peptides, and the known interaction with both USP30 (24) and with PINK1 (51-54) suggest that this may be a critical pUb nucleation site regulated by USP30 (Figure 7).

**Figure 7.**
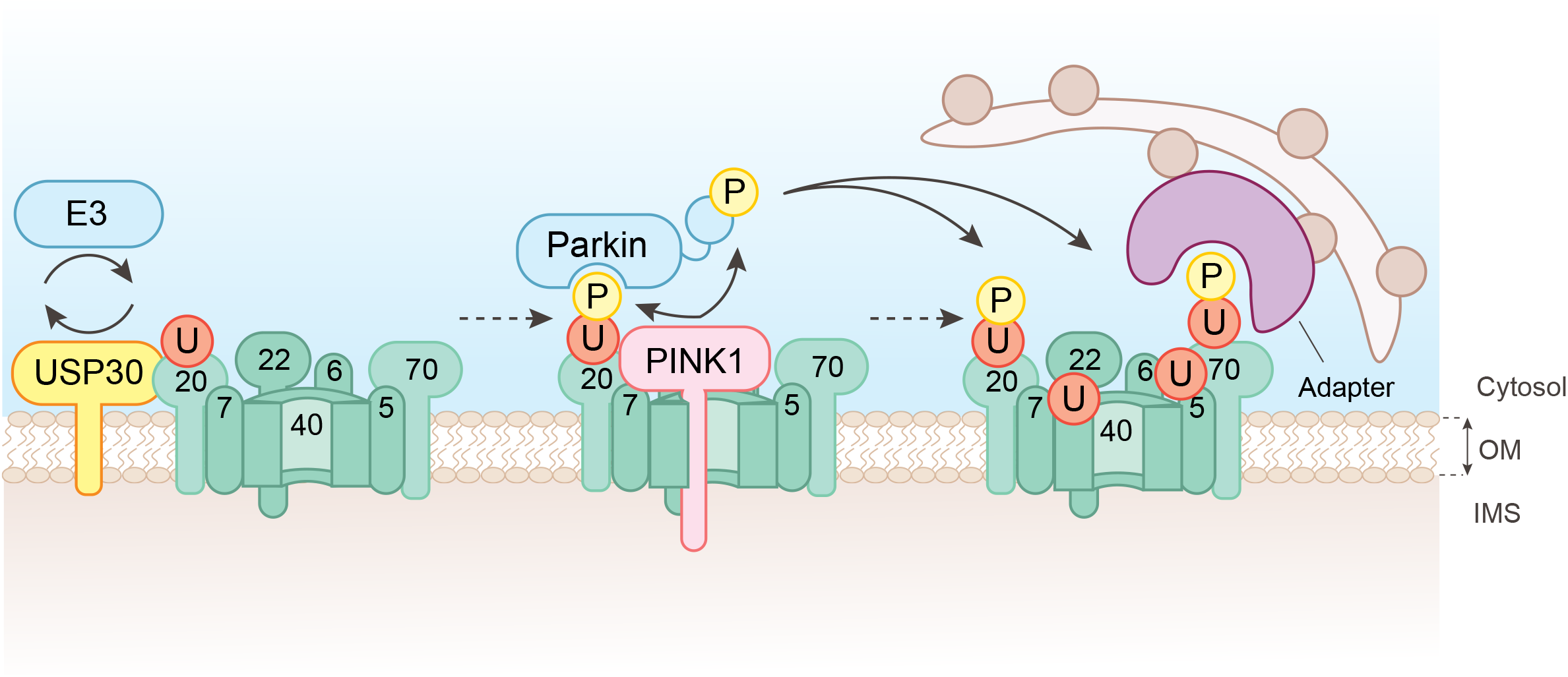
Working model depicting USP30 action upstream of PINK1. Under depolarising conditions PINK1 becomes activated but remains associated with TOM complex components. TOM complex associated ubiquitylation provides the nucleating substrate for PINK1-dependent phosphorylation of ubiquitin on Ser65. This leads to recruitment and activation of the E3 ligase Parkin, which can then amplify the signal. By opposing TOM complex ubiquitylation, USP30 suppresses the trigger for mitophagy.

Whilst our manuscript was in preparation, two complementary studies have been published that also highlight the centrality of the TOM complex to USP30 function (56, 57). All three studies use global proteome and ubiquitylome profiling. Ordureau et al. generate iNeurons and examine the impact of USP30 genetic loss (56). Phu et al. use HEK293 cells to compare genetic loss with a USP30 inhibitor that is related to the one we describe here (57). Note that, in that latter study a much higher concentration of inhibitor has been used (5μM vs 200nM). Both of those contemporaneous studies identify an overlapping set of USP30-sensitive ubiquitylation sites. They find greater prevalence, than we do here, of elevated ubiquitylation of mitochondrial matrix or inner mitochondrial membrane proteins, although we do see a few examples of the same phenomenon (e.g. MDH2, GRSF1, MTLN). Although ubiquitylation can occur within mitochondria (58), USP30 is an outer mitochondrial membrane protein whose catalytic activity is facing towards the cytosol (18, 20). Hence, it has been suggested that this reflects ubiquitylation of newly synthesised proteins engaging with the TOM complex (56, 57). Thus USP30 might sit at the gate of the import complex pore and strip off ubiquitin as a prerequisite for entry. This provides a striking parallel with the action of proteasomal deubiquitylases, which control entry to the proteasome core (59). Ribosomes themselves interact directly with the TOM complex (60) and ribosomal quality control (RQC) mechanisms have extensive links to the ubiquitin system (61). Perturbation of these pathways, could also lead to a higher representation of ubiquitylated peptides derived from nascent imported proteins. Our finding that the mitochondrial peptidyl-tRNA hydrolase PTRH2 is a USP30 substrate provides a first link to RQC. PTRH2 can cleave nascent chain tRNA on stalled ribosomes and provide a release mechanism for non-ubiquitylated nascent chains (50).

The USP30 dependent suppression of mitophagy is well established for events which rely on the over-expression of Parkin, together with acute mitochondrial depolarisation (22-24). In fact, in a recently published whole genome screen for mitophagy regulators in Parkin overexpressing C2C12 myoblasts, USP30 is the most prominent mitochondrial annotated negative regulator (25). Our study contributes to a body of evidence that translates these finding to systems with endogenous Parkin expression levels (20, 56, 57). The physiological defects associated with PINK1/Parkin loss of function in Parkinson’s Disease are likely to accumulate slowly. The benign effects of USP30 loss or inhibition, make it a target candidate that can be considered for long term therapy. The availability of specific tool compounds, such as described here, will enable pre-clinical assessment of this strategy.

## Acknowledgements

Funding for development of FT385 was provided by Forma Therapeutics. Additional support was provided by Celgene, Michael J. Fox Foundation, Alzheimers Research UK (MG, EM, KE), Parkinson’s UK (JJ), MRC (EM, AK, KM), Wellcome Trust (FGB, AG), European Union (FL). Funding for ODDI was obtained by John Davis. We thank Jon Lane and Ian Ganley for provision of cell lines.

## Methods

### Cell culture

hTERT-RPE1-YFP-PARKIN (24), SHSY5Y and SHSY5Y-mitoQC (mCherry-GFP-Fis1(101-152)) (42) cells were routinely cultured in Dulbecco’s Modified Eagle’s medium DMEM/F12 supplemented with 10% FBS and 1% non–essential amino acids.

### Generation of USP30 knockout cells

USP30 knockout cells were generated using CRISPR-Cas9 with USP30 specific sgRNAs targeting exon 3 of isoform 1 (sgRNA1: AGTTCACCTCCCAGTACTCC, sgRNA2: GTCTGCCTGTCCTGCTTTCA). sgRNAs were cloned into the pSpCas9(BB)–2A–GFP (PX458) vector (Addgene plasmid #48138 46) or PX330-Puro (kind gift from Prof Ciaran Morrison, NUI Galway). hTERT–RPE1–YFP-Parkin USP30 knockout Clone 1E and SHSY5Y clone D were engineered by transfecting the parental lines with pSpCas9(BB)–2A–GFP-sgRNA1, followed by FACS 24 hours later (selection for GFP positive cells) and single cell dilution. SHSY5Y-mitoQC Clone 11 was engineered by transfection with PX330-Puro-sgRNA2 followed by selection with 1-1.5 μg/ml Puromycin and single cell dilution. Individual clones of SHSY5Y KO cells were amplified and multiple alleles sequenced (Supplementary Figure 1). The positive clone (KO11) obtained has lost expression of the mitoQC fluorophore.

### Antibodies and reagents

Antibodies and other reagents used were as follows: anti–USP30 (Sigma HPA016952, 1:500), anti-USP30 (Thermo Fisher, PA5-53523, 1:1000), anti-USP30 (MRC PPU, Dundee, 1:1000), anti-USP30 (Santa-Cruz, sc-515235, 1:1000), anti–PINK1 (Fig 1E; D8G3, Cell Signalling Technology, 6946S, 1:1000), anti-TOM20 (Sigma HPA011562, 1:1,000), anti-PARK2 (SantaCruz, sc32282, 1:250), anti-MFN2 (Abcam, ab56889, 1:1000), anti-ubiquitin (Lifesensor, VU101, 1:2000), anti-Fis1 (ProteinTech, 10956-1-AP, 1:1000), anti-phospho-Ubiquitin Ser65 (Millipore, ABS1513-I, 1:1000), anti-phosphoUbiquitin Ser65 (Cell Signalling Technology, 62802, 1:1000), anti–VDAC1 (AbCam, ab15895, 1:1000), mouse anti–actin (AbCam ab6276, 1:10,000), mouse anti-actin (ProteinTech 66009-1-Ig, 1:10,000), rabbit-anti-actin (ProteinTech,20536-1-AP, 1:10,000), anti-SYNJ2BP (Sigma HPA000866, 1:1000), oligomycin A (SIGMA 75351), antimycin A (SIGMA A8674).

### Preparation cell lysates and Western blot analysis

Cultured cells were either lysed with urea buffer (Figure 6E, 9M urea, 20mM Hepes-NaOH pH7.4) supplemented with 2-Chloroacetamide (CAA, Sigma) or NP–40 (0.5% NP–40, 25 mM Tris-HCl pH 7.5, 100 mM NaCl, 50 mM NaF) lysis buffer and routinely supplemented with mammalian protease inhibitor (MPI) cocktail (Sigma) and Phostop (Roche), with the exception of data presented in Figure 2. Proteins were resolved using SDS–PAGE (Invitrogen NuPage gel 4–12%), transferred to nitrocellulose membrane, blocked in 5% milk, 5% BSA or 0.1% Fish Skin Gelatin in TBS supplemented with Tween–20, and probed with primary antibodies overnight. Visualisation and quantification of Western blots were performed using IRdye 800CW and 680LT coupled secondary antibodies and an Odyssey infrared scanner (LI–COR Biosciences, Lincoln, NE).

### Sub-cellular fractionation

SHSY5Y cells were washed with ice-cold PBS and then collected by scraping and centrifugation at 1000g for 2mins. Cell pellets were washed with HIM buffer (200 mM mannitol, 70 mM sucrose, 1 mM EGTA, 10 mM HEPES-NaOH pH 7.4) and then resuspended in HIM buffer supplemented with mammalian protease inhibitors. Cells were mechanically disrupted by shearing through a syringe with a 27G needle, followed by passing 3 times though a 8.02mm diameter “cell cracker” homogeniser using a 8.01mm diameter ball bearing (62) or passage through a 27G needle (Figure 2A). The resulting homogenate was cleared from nuclei and unbroken cells by centrifugation at 600g for 10 minutes to obtain a post nuclear supernatant (PNS). The PNS was separated into the post-mitochondrial supernatant (PMS) and crude mitochondrial fraction (MF) by centrifugation at 7000g for 15mins. The MF pellet was resuspended in HIM buffer + MPI.

### Activity probe assay

Cells were mechanically homogenised in HIM buffer supplemented with 1mM DTT (Figure 2A) or 1mM Tris(2-carboxyethyl)phosphine (TCEP, Figure 2B) to obtain the PNS. Homogenates were incubated with Ub-propargyl (Ub-PA, UbiQ) probe at 1:100 (w/w) for 15 minutes at 37°C (40). The reaction was stopped by the addition of sample buffer and heating at 95°C. To test drug engagement either intact cells or cell homogenate (PNS, without addition of protease inhibitors) were treated with FT385. Intact cells were treated for 4hrs at 37°C prior to homogenisation and homogenate was pre-incubated for 30mins at room temperature before probe incubation. Samples were either processed using a WES system and transformed to a virtual Western blot (Figure 2A, Protein Simple, Biotechne) or analysed by standard Western blot (Figure 2B).

### SILAC Labelling

SHSY5Y and SHSY5Y-KO11 cells were grown for at least 8 passages in SILAC DMEM/F12 supplemented with 10% dialysed FBS, 200 mg/L L-proline and either L-lysine (Lys0) together with L-arginine (Arg0), L-lysine-^2^H_4_ (Lys4) with L-arginine-U-^13^C_6_ (Arg6) or L-lysine-U-^13^C_6_-^15^N2 (Lys8) with L-arginine-U-^13^C_6_-^15^N_4_ (Arg10) at final concentrations of 28 mg/L arginine and 146 mg/L lysine.

### Proteomics methods

For the experiments shown in Figure 4, SILAC labelled cells, were lysed by sonication in 9M urea, 20mM HEPES pH 8.0, 1mM sodium orthovanadate, 2.5mM sodium pyrophosphate, 1mM glycerol-3-phosphate. For total proteome and ubiquitylome, 700μg and 20mg respectively, of each sample was combined at a 1:1:1 ratio respectively. For the experiments shown in Figure 5, Mitochondrial fractions (ubiquitylome) were obtained by homogenisation in HIM buffer supplemented with mammalian protease inhibitors, 50mM CAA and Phostop from SILAC labelled cells. Cell pellets (proteome) or MFs were lysed by sonication in 9M urea, 20 mM HEPES pH 8.0, 1.15 mM sodium molybdate, 1mM sodium orthovanadate, 4 mM sodium tartrate dihydrate, 5mM glycerol-3-phosphate, 1mM sodium fluoride, then reduced and alkylated with either 4.5mM dithiothreitol/10mM iodoacetamide (Figure 4) or 10mM TCEP/10mM CAA (Figure 5). Urea was then diluted 4 fold by the addition of 20mM HEPES buffer prior to trypsinisation overnight. The resultant tryptic peptides were acidified with Trifluoroacetic acid and purified on a C18 Sep-Pak column before lyophilisation (Figure 4) or drying with a SpeedVac (Figure 5).

For ubiquitylome samples, modified peptides were enriched by immunoprecipitation using a diGly specific antibody in accordance with manufacturer’s instructions (PTMScan Ubiquitin Remnant Motif (K-GG) Kit #5562, Cell Signaling Technology). Eluted peptides were purified using C18 stage tips (Figure 4) or C18 Sep-Pak columns (Figure 5). Samples were then dried in a speed-vac before resuspension and analysis by LC-MS/MS. Ubiquitylome (Figure 4) samples were analysed (total 5 technical replicates) on an Orbitrap Fusion Lumos (1 replicate) and Orbitrap Q Exactive HF (4 replicates). Ubiquitylome (Figure 5) samples were analysed on an Orbitrap Fusion Lumos.

For proteome samples, peptides were separated by fractionation. For Figure 4, samples were fractionated by off-line high-pH reverse-phase pre-fractionation as previously described (63), with the exception that eluted peptides were concatenated down to 10 fractions. Briefly, digested material was fractionated using the loading pump of a Dionex Ultimate 3000 HPLC with an automated fraction collector and a XBridge BEH C18 XP column (3 × 150 mm, 2.5μm particle size, Waters no. 186006710) over a 100 min gradient using basic pH reverse-phase buffers (A: water, pH 10 with ammonium hydroxide; B: 90% acetonitrile, pH 10 with ammonium hydroxide). The gradient consisted of a 12 minute wash with 1% B, then increasing to 35% B over 60 minutes, with a further increase to 95% B in 8 minutes, followed by a 10 minute wash at 95% B and a 10 minute re-equilibration at 1% B, all at a flow rate of 200μl/min with fractions collected every 2 minutes throughout the run. 100μl of the fractions was dried and resuspended in 20μL of 2% acetonitrile/0.1% formic acid for analysis by LC-MS/MS. Fractions were loaded on the LC-MS/MS following concatenation of 50 fractions into 10, combining fractions in a 10-fraction interval (F1 + F11 + F21 + F31 + F41… to F10 + F20 + F30 + F40 + F50). For Figure 5, samples were fractionated by off-line reverse-phase prefractionation using a Dionex Ultimate 3000 Off-line LC system. Briefly, digested material was fractionated using the loading pump of a Dionex Ultimate 3000 HPLC with an automated fraction collector and with a Gemini C18, (3um particle size, 110A pore, 3 mm internal diameter, 250 mm length, Phenomenex #00G-4439-Y) over a 39 minute gradient using the following buffers: A: 20mM Ammonium Formate, pH=8; B: 100% ACN. The gradient consisted of a 1 minute wash with 1% B, then increasing to 35.7% B over 28 minutes,< followed by a 5 minute wash at 90% B and a 5 minute re-equilibration at 1% B, all at a flow rate of 250 μl/min. with fractions collected every 45s from 2 minutes to 38 minutes for a total of 48 fractions. Non-consecutive concatenation of every 13th fraction was used to obtain 12 pooled fractions (Pooled Fraction 1: Fraction 1 + 13 + 25 + 27, Pooled Fraction 2: Fraction 2 + 14 + 26 + 38 …).

### Orbitrap Q Exactive HF LC-MS/MS parameters

Peptide fractions were analysed by nano-UPLC-MS/MS using a Dionex Ultimate 3000 nano UPLC with EASY spray column (75μm x 500 mm, 2μm particle size, Thermo Scientific) with a 60 minute gradient of 2% acetonitrile, 0.1% formic acid in 5% DMSO to 35% acetonitrile, 0.1% formic acid in 5% DMSO at a flow rate of ~250nl/minute. MS data was acquired with an Orbitrap Q Exactive HF instrument in which survey scans were acquired at a resolution of 60.000 @ 200m/z and the 20 most abundant precursors were selected for HCD fragmentation with a normalised collision energy of 28.

### Orbitrap Fusion Lumos LC-MS/MS parameters

Samples (Figures 4 & 5) were analysed by LC-MS/MS on a Dionex Ultimate 3000 connected to an Orbitrap Fusion Lumos. For experiments presented in Figure 4 peptides were separated using a 60 minute linear gradient from 2–35 % acetonitrile in 5% DMSO, 0.1% formic acid at a flow rate of 250nl/min. on a 50cm EASY spray column (75μm x 500mm, 2μm particle size, Thermo Scientific). For experiments presented in Figure 5 peptides were separated using 120 (proteome) or 240 (ubiquitylome) minute linear gradients from 0–28 % acetonitrile in 3% DMSO, 0.1% formic acid at a flow rate of 300nl/minute on a 50cm EASY spray column (75μm x 500mm, 2μm particle size, Thermo Scientific). MS1 scans were acquired at a resolution of 120,000 between 400 – 1,500 m/z with an AGC target of 4×10^5^. Selected precursors were fragmented using HCD at a normalised collision energy of 28% (Figure 4) or 30% (Figure 5), an AGC target of 4×10^3^ (Figure 4) or 4 ×10^4^ (Figure 5), a maximum injection time of 35ms (Figure 4) or 45ms (Figure 5), a maximum duty cycle of 1 second (Figure 4) or 3 seconds (Figure 5) and a dynamic exclusion window of 60 seconds (Figure 4) or 45seconds (Figure 5). MS/MS spectra were acquired in the ion trap using the rapid scan mode.

### Mass spectrometry data analysis

All raw MS files from the biological replicates of the SILAC-proteome experiments were processed with the MaxQuant software suite; version 1.6.7.0 using the Uniprot database (retrieved in July 2019) and the default settings (64). Cysteine carbamidomethylation was set as a fixed modification, whereas oxidation, phospho(STY), GlyGly (K) and acetyl N terminal were considered as variable modifications. Data was requantified. ProteinGroup text files (proteome) or GlyGly (K) sites files were further processed using Excel (see Supplementary Table 1) and Perseus (version 1.6.10.50). Graphs were plotted using JMP13.

### In Vitro USP30 Activity Assay

Fluorescence intensity measurements were used to monitor the cleavage of a ubiquitin-rhodamine substrate. All activity assays were performed in black 384-well plates in 20 mM Tris-HCl, pH 8.0, 0.01 % Triton-X, 1 mM L-Glutathione and 0.03% Bovine Gamma Globulin with a final assay volume of 20μl. Compound IC50 values for DUB inhibition were determined as previously described (33). Briefly, an 11-point dilution series of compounds were dispensed into black 384-well plates using an Echo (Labcyte). USP30, 0.2 nM (#E-582 residues 57-517, Boston Biochem) was added and the plates pre-incubated for 30 minutes. 25 nM ubiquitin-rhodamine 110 (Ubiquigent) was added to initiate the reaction and the fluorescence intensity was recorded for 30 minutes on a Pherastar FSX (BMG Labtech) with a 485 nm excitation/520 nm emission optic module. Initial rates were plotted against compound concentration to determine IC_50_.

### k_*inact*_/K_*I*_ determination

A k_inact_/K_I_ assay was carried out using 0.2nM USP30 and 180nM ubiquitin-rhodamine 110 as described above with the omission of the 30 minute pre-incubation step. Upon addition of the substrate, fluorescence intensity was monitored kinetically over 30 minutes. Analysis was performed in GraphPad Prism. Kinetic progress curves were fitted to equation y = (v*i*/k_obs_)(1−exp(−k_obs_x)) to determine the k_obs_ value. The k_obs_ value was then plotted against the inhibitor concentration and fitted to the equation y = k_inact_/(1 + (K_I_/x)) to determine k_inact_ and K_I_ values.

### Bio-layer interferometry

Bio-layer interferometry was performed on an Octet RED384^®^ system (ForteBio) at 25°C in a buffer containing 50mM HEPES buffer (pH 7.5), 400mM NaCl, 2mM TCEP, 0.1% Tween, 5% Glycerol and 2% DMSO. Biotinylated USP30 (residues 64-502Δ179-216 & 288-305, Viva Biotech Ltd., Shanghai) was loaded onto Superstreptavidin (SSA) biosensors. Association of defined concentrations of FT3967385 (0 – 6.67μM) was recorded over 180 seconds followed by dissociation in buffer over 600 seconds. Traces were normalised by double subtraction of baseline (no USP30, no compound) and reference sensors (no USP30, association and dissociation of compound) to correct for non-specific binding to the sensors. Traces were analysed using Octet Software (Version 11.2, ForteBio).

### Live cell imaging and basal mitophagy quantification

SHSY5Y cells stably expressing mCherry-GFP-Fis1(101-152) (SHSY5Y mitoQC) were treated every 24 hours over a 96 hrs timecourse with 200 and 500nM of FT385. Cells were replated onto a IBIDI μ-Dish (2×10^5^) two days before live-cell imaging with a 3i Marianas spinning disk confocal microscope (63x oil objective, NA 1.4, Photometrics Evolve EMCCD camera, Slide Book 3i v3.0). Cells were randomly selected using the GFP signal and images acquired sequentially (488nm laser, 525/30 emission; 561nm laser, 617/73 emission). Analysis of mitophagy levels was performed using the ‘mito-QC Counter’ implemented in FIJI v2.0 software (ImageJ, NIH) as previously described (65), using the following parameters: Radius for smoothing images = 1.25, Ratio threshold = 0.8, and Red channel threshold = mean + 0.5 standard deviation. Mitophagy analysis was performed for three independent experiments with 80 cells per condition. One-Way ANOVAs with Dunnett’s multiple comparisons were performed using GraphPad Prism 6. *P*-values are represented as \*\**P* < 0.01, \*\*\*\**P* < 0.0001. Error bars denote standard deviation.

## Supplementary Figure legends

**Supplementary Figure 1.**
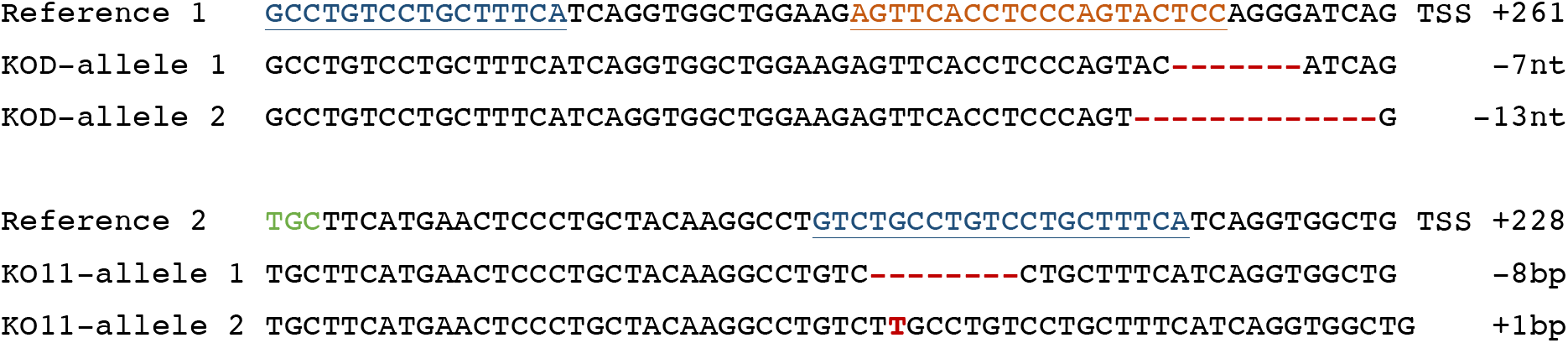
sgRNA design for the generation of USP30 KO cell lines and sequence of the two SHSY5Y USP30 KO clones used in this study. KOD was generated using sgRNA#1 (target region shown in orange) and KO11 was generated using sgRNA#2 (target region shown in blue). Insertions and deletions are indicated in red. TSS: transcriptional start site. The green codon corresponds to catalytic C77. Frequency of allele detection: KO11 allele 1 (4/5), allele 2 (1/5); KOD allele 1 (7/10), allele 2 (3/10).

**Supplementary Figure 2.**
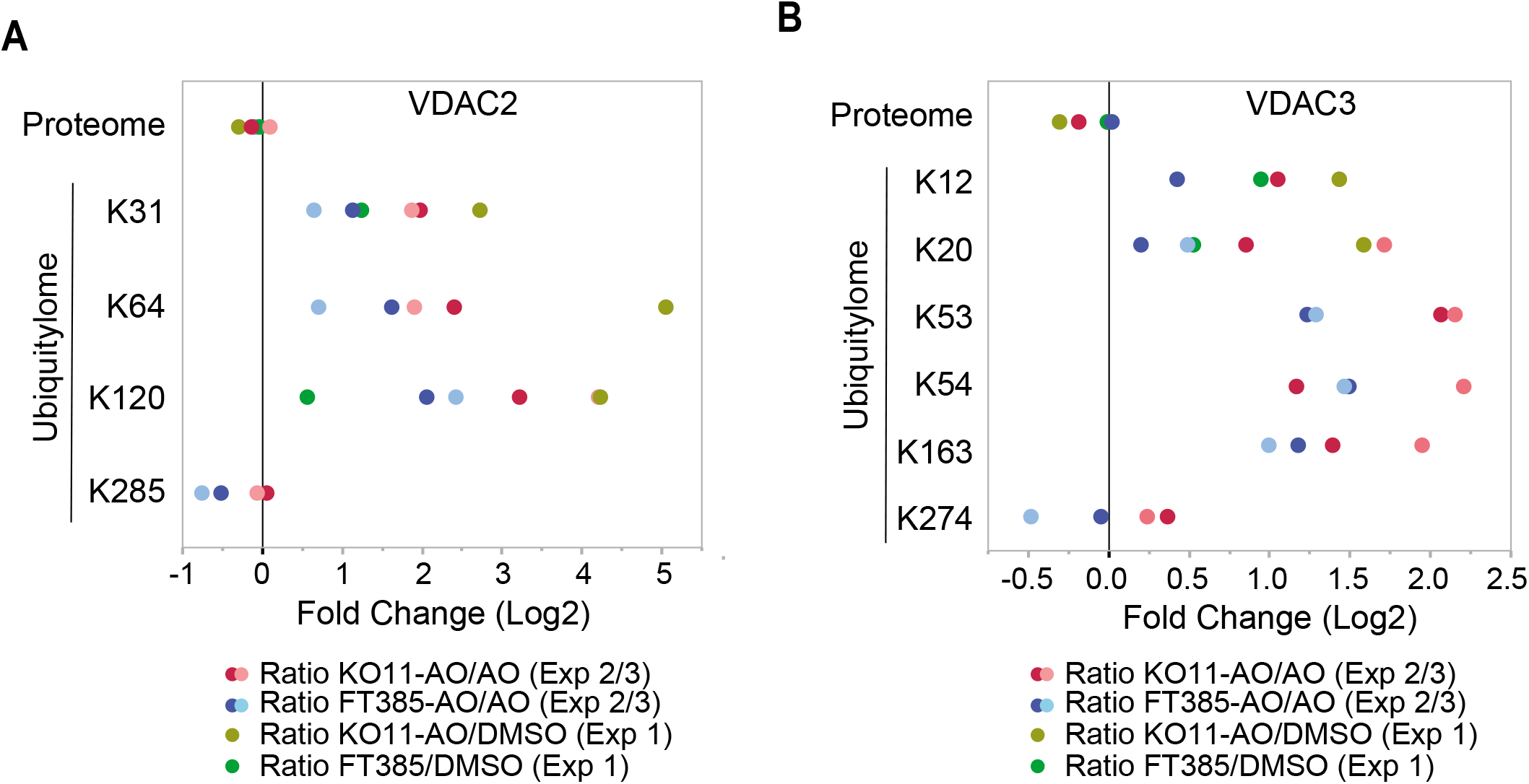
Comparison of VDAC2 and VDAC3 Ubiquitylome changes in USP30 KO versus USP30 inhibitor treated SHSY5Y cells. Fold change (Log2) in proteome and individual diGly peptides (localisation ≥0.75) by site in VDAC2 and VDAC3 proteins (right). Complements Figure 4E.

**Supplementary Figure 3.**
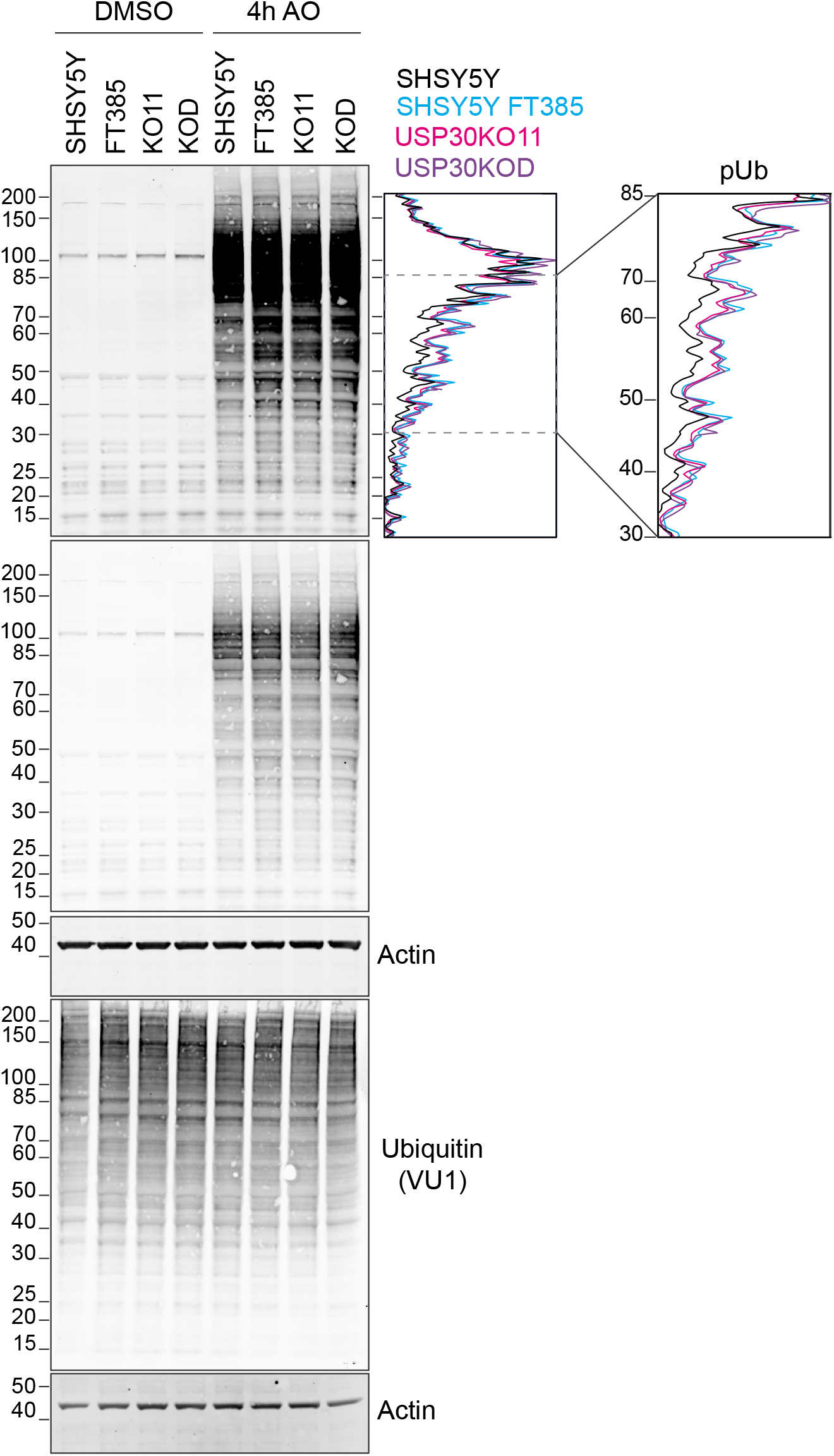
Comparison of depolarisation induced phosphoSer65-Ubiquitin (pUb) generation in SHSY5Y cells treated with FT385 and USP30 KO SHSY5Y (KO11 and KOD). Shown is a western blot of the same samples shown in Figure 3C, and corresponding line-graph for the pUb signal, of lysates from cells treated for 4 hours with AO (1μM) with or without FT385 (200 nM). Red arrows indicate ubiquitylated TOM20 and the arrowheads indicate unmodified TOM20.

## Notes

### Competing Interest Statement

M.G. is receiving research funding from AstraZeneca, Pfizer and Merck
MC,SU,HM and ODDI have received research funding from Celgene.

